# Noise-Transfer2Clean: Denoising cryo-EM images based on noise modeling and transfer

**DOI:** 10.1101/2021.05.10.443396

**Authors:** Hongjia Li, Hui Zhang, Xiaohua Wan, Zhidong Yang, Chengmin Li, Jintao Li, Renmin Han, Ping Zhu, Fa Zhang

## Abstract

**Motivation:** Cryo-electron microscopy (cryo-EM) is a widely-used technology for ultrastructure determination, which constructs the three-dimensional (3D) structures of protein and macromolecular complex from a set of two-dimensional (2D) micrographs. However, limited by the electron beam dose, the micrographs in cryo-EM generally suffer from extremely low signal-to-noise ratio (SNR), which hampers the efficiency and effectiveness of downstream analysis. Especially, the noise in cryo-EM is not simple additive or multiplicative noise whose statistical characteristics are quite different from the ones in natural image, extremely shackling the performance of conventional denoising methods.

**Results:** Here, we introduce the Noise-Transfer2Clean (NT2C), a denoising deep neural network (DNN) for cryo-EM to enhance image contrast and restore specimen signal, whose main idea is to improve the denoising performance by correctly discovering the noise model of cryo-EM images and transferring the statistical nature of noise into the denoiser. Especially, to cope with the complex noise model in cryo-EM, we design a contrast-guided noise and signal re-weighted algorithm to achieve clean-noisy data synthesis and data augmentation, making our method authentically achieve signal restoration based on noise’s true properties. To our knowledge, NT2C is the first denoising method that resolves the complex noise model in cryo-EM images. Comprehensive experimental results on simulated datasets and real datasets show that NT2C achieved a notable improvement in image denoising and specimen signal restoration, comparing with the state-of-art methods. A real-world case study shows that NT2C can improve the recognition rate on hard-to-identify particles by 19% in the particle picking task.

## 1 INTRODUCTION

Cryo-electron microscopy (cryo-EM) is a widely-used technology that resolves high-resolution three-dimensional (3D) structures of protein and macromolecular complexes from a series of two-dimensional (2D) micrographs (Bai *et al*., 2015). However, the signal-to-noise ratio (SNR) of raw cryo-EM images is estimated to be only as high as 0.01∼0.1 (Bendory *et al*., 2020), amongst the lowest in any imaging field, which extremely decreases the accuracy and efficiency in downstream analysis of cryo-EM images and reduces the confidence of structures determination. Therefore, an image restoration operation is usually necessary before particle picking, structure segmentation, and other cryo-EM data analysis processes to attain high-resolution cryo-EM 3D reconstructions.

A variety of conventional methods have been developed to improve the contrast and decrease the noise level in cryo-EM micrographs, such as BM3D (Dabov *et al*., 2007), band-pass filter (Penczek *et al*., 2010) and Wiener filter (Sindelar *et al*., 2011). For an image restoration algorithm, additional image prior knowledge will be introduced to repair the missing and degenerated information, in which human knowledge concluded from the natural images are usually used. However, the noise model in cryo-EM micrograph is usually unknown and varies in different data collection configurations. Therefore, the pre-defined image priors used in these conventional methods cannot correctly fit the noise model in cryo-EM, leading to a limited performance when the conventional methods are applied to cryo-EM data.

Recently, learning-based denoising methods have shown their advantages. Mao *et al*., 2016 proposed an encoding-decoding framework with symmetric convolutional-deconvolutional layers for image restoration. Ledig *et al*., 2017 presented a generative adversarial network (GAN) for image super-resolution (SR) which recovers photo-realistic textures from heavily downsampled images. However, most of these learning methods require a clean-noisy paired dataset for training, therefore, can not be applied to cryo-EM, where ground truth is unavailable. To overcome this barrier, several methods learned from paired noisy images or single noisy images are proposed (Krull *et al*., 2019). Lehtinen *et al*., 2018 presented a general machine learning (ML) framework, called Noise2Noise (N2N), for learning denoising models from paired noisy images. Chen *et al*., 2018 proposed a GAN-CNN based framework, GCBD, for learning denoising models from single noisy images where GAN (Goodfellow *et al*., 2012) is utilized to build paired training datasets and then convolutional neural network (CNN) is employed for denoising. Specifically, Bepler *et al*., 2020 proposed a denoiser called Topaz-Denoise for cryo-EM and cryo-ET, based on an N2N architecture and trained by thousands of cryo-EM micrographs.

In this paper, we propose a novel denoising framework, the Noise-Transfer2Clean (NT2C), to restore the specimen signal and enhance the image contrast by discovering the unknown noise model from the cryo-EM image. Firstly, a coarse CNN denoiser is trained to enhance the contrast of a cryo-EM image, to distinguish the background and specimen signal. Then, the pure noise patches are extracted from the micrographs and fed into a GAN to estimate and simulate the noise distribution. Finally, a fine denoising network is able to be trained by the clean-noisy pairs simulated from the accurately estimated noise distribution in GAN. By resolving the noise model in a cryo-EM image, our strategy is able to further decomplex the specimen signal from the noisy background. Especially, to cope with the complex noise model in cryo-EM, we design a contrast-guided noise and signal re-weighted algorithm to achieve clean-noisy data synthesis and data augmentation, making our method authentically achieve signal restoration based on the noise’s true properties. We have tested and compared our denoising model with several commonly-used cryo-EM denoising algorithms on both synthetic and real datasets. The experiment results on synthetic datasets show NT2C’s denoising performance is comparable to current state-of-the-art methods; and the results on three real datasets demonstrate NT2C’s ability to deal with real-world high-noise-level cryo-EM images. A case study on particle picking further proves that our denoising method is able to restore very weak particle signals, improving the recognition rate on hard-to-identify particles by 19%.

## 2 THE IMAGING MODEL IN CRYO-EM

The imaging process in cryo-EM involves the conversion of the electron wave’s intensity distribution into a digital signal via a detector (Vulović *et al*., 2013), which introduces multiple complex noises to the final micrographs. According to the principle of cryo-EM, the detected image can be written as

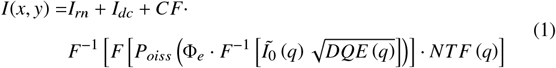

where 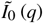 is the Fourier spectrum of the noise-free signal, *DQE* is the detective quantum efficiency, Φ_*e*_ is the incident electron flux in [*e*^−^/*area*], *F* is Fourier transform, *F*^−1^ is inverse Fourier transform, *P_oiss_* is the Poisson described shot noise, *NTF* is the noise transfer function, *CF* is a scale factor for detectors, *I_rn_* indicates the readout noise and *I_dc_* indicates the dark current of detector. The detective quantum efficiency *DQE* is defined as *DQE* (*q*) = *MTF*^2^ (*q*)/*NTF*^2^ (*q*), where *NTF*^2^ (*q*) = *NPS_out_*/(*CF*^2^Φ_*e*_ with *NPS* being the noise power spectrum and *MTF* describes transfer of the signal amplitude for different spatial frequencies. As described in the formula, the signal propagation can be modeled as follows: (1) 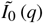 (*q*) is damped (multiplied) by the ratio between signal (*MTF*) and noise (*NTF*) transfer, (2) the signal is multiplied by the integrated electron flux and shot noise contributions are added, (3) the Fourier spectrum of that noisy signal is damped by the *NTF*, (4) the number of electrons are scaled to the image gray values in detector analog-to-digital unit [*ADU*]. Therefore, the noise in cryo-EM comes from three aspects: the shot noise contributions added in the measurement process; the readout noise *I_rn_* and dark current *I_dc_* from the detector; blurring effect caused by detector point spread function PSF (x,y) whose Fourier transform is the MTF.

## 3 METHODS

### 3.1 NT2C protocol

#### 3.1.1 Overview of the procedure

The key idea of NT2C is discovering the noise model of cryo-EM images over pure noise patches and transferring the statistical nature of noise into the denoiser. As shown in Figure 1, NT2C contains three modules:

**Fig. 1.**
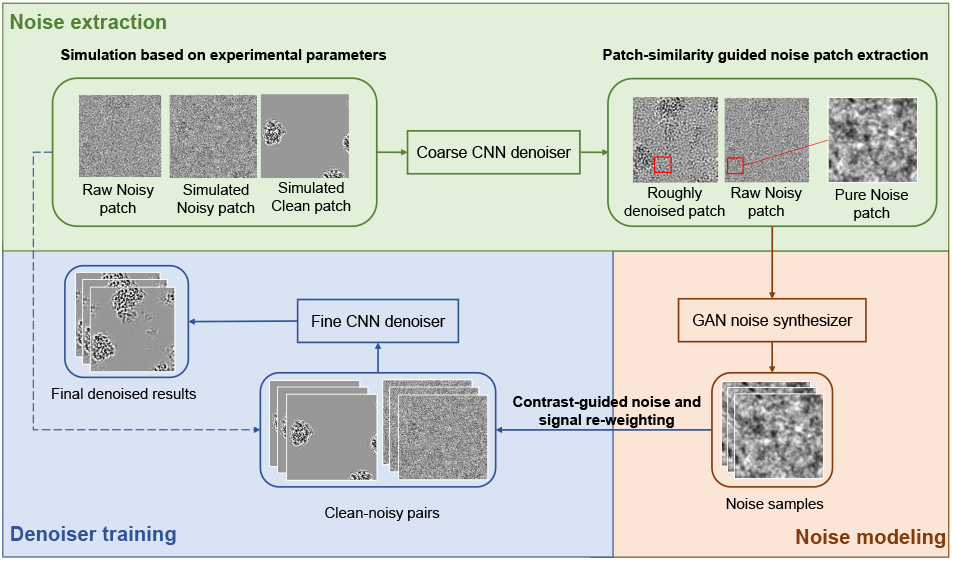
The overall protocol of NT2C method for cryo-EM image denoising. There are three main modules in NT2C: (a) noise extraction, (b) noise modeling and (c) denoiser training.

##### (a) Noise extraction

The noise extraction module takes raw micrographs as input, and output the pure noise patches of the background (see subsubsection 3.2.2). Due to the extremely low SNR in cryo-EM micrographs, distinguishing the background from the particles is a hard task in raw noisy micrographs. Here, a coarse denoiser (see subsubsection 3.1.2) is trained to roughly enhance the image quality and aid the extraction of noise patches, based on the simulated datasets with the same experimental parameters (see subsubsection 3.2.1).

##### (b) Noise modeling

The statistical properties of noise in cryo-EM micrograph change with different configurations during data collection. It is critical to correctly understand the statistical nature of the noise for image denoising. Here, a GAN noise synthesizer (see subsubsection 3.1.2) is trained to learn the statistical properties of noise, with pure noise patches as input and simulated noise patches as output.

##### (c) Denoiser training

The noise synthesizer poses the possibility of clean-noisy pair generation for cryo-EM, which is critical in denoiser training. However, as described in section 2, the noise pattern in cryo-EM is quite complex. Here, we design a contrast-guided noise and signal re-weighted algorithm (see subsubsection 3.2.3) to transfer the non-additive noise to a clean image, to achieve clean-noisy pair synthesis and data augmentation. Based on the abundant synthesized clean-noisy pairs, a fine denoiser (see subsubsection 3.1.2) is able to be trained to precisely restore specimen signal from the high-level noise.

#### 3.1.2 Main components

##### 1) CNN denoiser

The CNN denoiser in NT2C is based on a U-net architecture (Ronneberger *et al*., 2015), which contains five max-pooling down-sampling blocks and five nearest-neighbor up-sampling blocks, with skip connections between down- and up-sampling blocks at each spatial resolution (shown in Figure 2). Given the set of clean-noisy pairs {*y* and *x* ∼ *Noise* (*y*)}, a denoising function *f* with parameter *θ* can be learned. The loss function for our task is

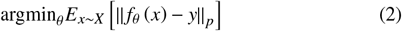

where *p* = 2 is used in NT2C to find *f* with mean-seeking behavior.

The CNN denoiser has been called twice in NT2C’s procedure: (i) as a coarse denoiser trained with the simulated clean-noisy pairs (see subsubsection 3.2.1) to roughly enhance the contrast of cryo-EM images, for the ease of noise patch extraction; (ii) as a fine denoiser trained with the clean-noisy pairs produced by the noise and signal re-weighted algorithm (see GAN noise synthesizer and subsubsection 3.2.3) to capture the nature of noise statistical properties and restore the specimen signal in cryo-EM micrographs. Here, the model parameters of the coarse denoiser could be transferred to the fine denoiser to avoid retrain from scratch.

**Fig. 2.**
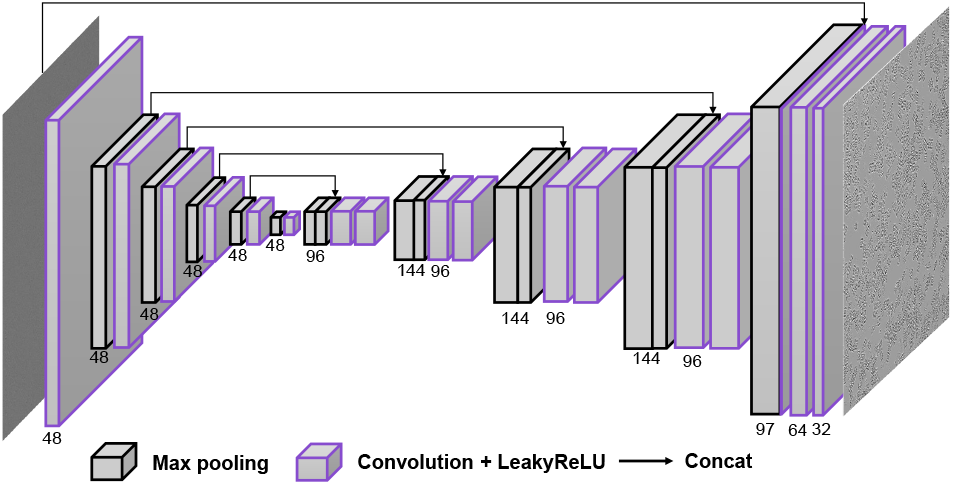
The network architecture of the CNN denoiser. The U-net model consists of 5 convolutional and down-sampling blocks followed by 5 convolutional and up-sampling blocks. Skip connections link each down-sampling block to the mirrored up-sampling block.

##### 2) GAN noise synthesizer

The noise model in cryo-EM is too complex to be explicitly described by an analytical expression. Here, the improved GAN framework (Gulrajani *et al*., 2017) is adopted to implicitly learn the latent noise model in cryo-EM micrographs. This framework contains two components, a generative network that consists of five fully connected layers, and a discriminative network which consists of three full connected layers (shown in Figure 3). Batch Normalization (Ioffe *et al*., 2015) is used in the generator to secure the model stability and Leaky Rectified Linear Unit (LeakyReLU) (Maas *et al*., 2013) activation is used in both the generator and the discriminator to ensure fast learning.

**Fig. 3.**
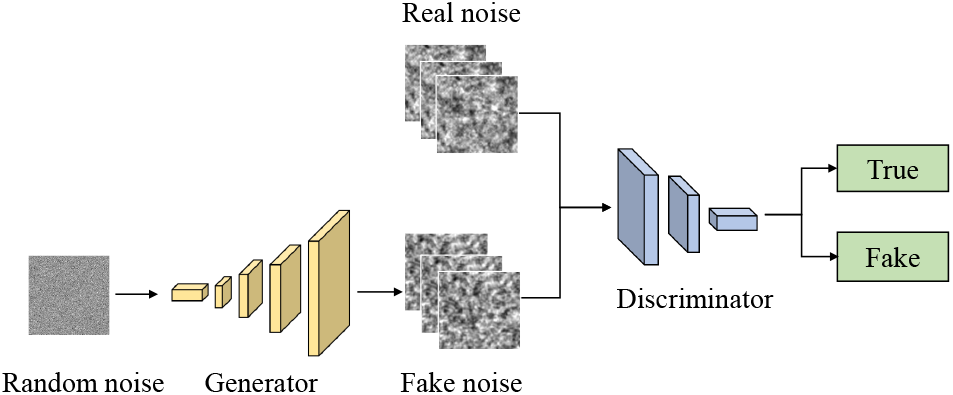
The network architecture of the GAN noise synthesizer. The generative network is trained to generate noise samples, while the discriminative network is trained to determine whether a noise sample is from real data or the generative network.

The generative network is trained to generate noise samples while the discriminative network is trained to determine whether a sample is from real data or the generative network. After the convergence of adversarial learning, the generative network will be able to produce noise patches hard to be distinguished from real noise patches. The loss function for our task is

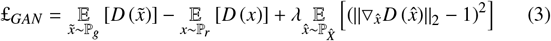

where ℙ_*r*_ is the distribution over noise patches, ℙ_*g*_ is the generator distribution, 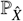 is defined as a distribution sampling uniformly along straight lines between pairs of points sampled from P_*r*_ and P_*g*_.

### 3.2 Detailed algorithm

#### 3.2.1 Simulation based on experimental parameters

The lack of supervised training data hampers the application of learning-based denoising method in cryo-EM. Here, we adopt the simulation software InSilicoTEM (Vulović *et al*., 2013) to generate paired clean-noisy datasets, which can simulate the photographing process in cryo-EM based on physical principles.

We set the simulation according to the experimental parameters used in data collection (shown in Figure 4), including the pixel size, defocus, voltage, electron dose, and detector type, which make the limited resolution, contrast transfer function (Wade *et al*., 1992) and modulation in the simulation very close to the real-world data. The homologous proteins with similar size downloaded from Protein Data Bank (PDB) (Burley *et al*., 2017) are used to produce clean ground truth. Such a simulation is possible to generate datasets with statistical properties of noise close to the ones in real cryo-EM micrographs. Consequently, the composed clean-noisy pairs could be fed into the coarse CNN denoiser to produce a model that achieves roughly denoising and contrast enhancement on the cryo-EM image, for the ease of noise patch extraction.

**Fig. 4.**
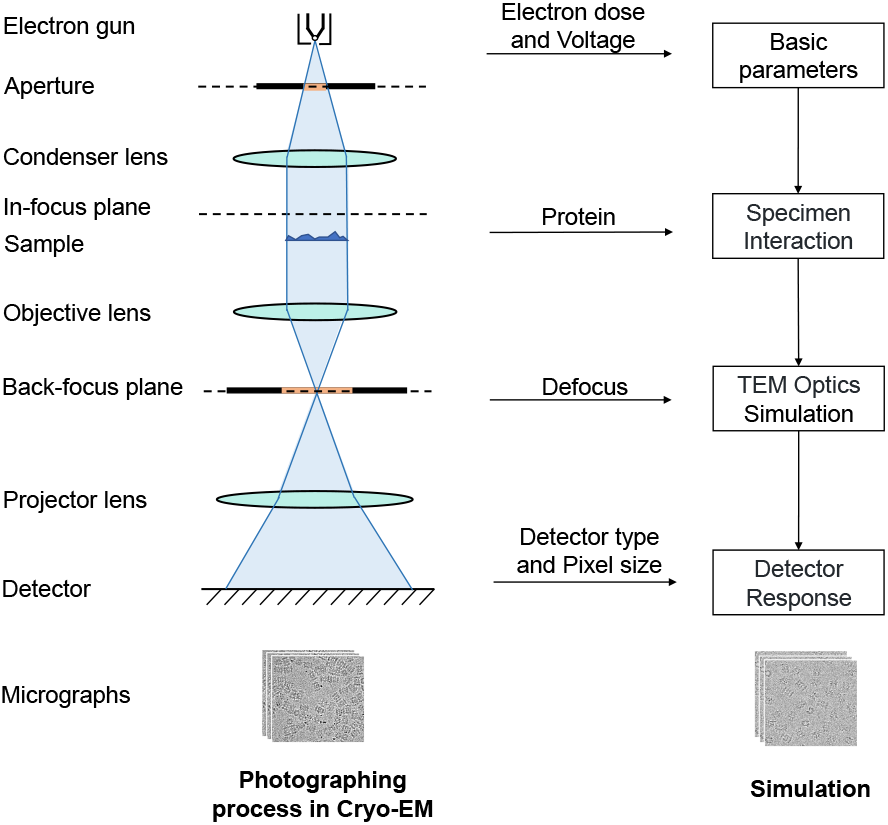
Simulation based on experimental parameters. The experimental parameters in photographing process (left) are used for simulation (right), including electron dose, voltage, defocus, pixel size, and detector type. The homologous protein with similar size is utilized for sample simulation.

#### 3.2.2 Patch-similarity guided noise patch extraction

If we divide a cryo-EM micrograph into patches with suitable size, these patches could be classified into two categories: patch containing specimen signal or patch of background with pure noise (shown in Figure 5). Naturally, the background patches are homogeneous to each other while the patches containing specimen signals have different patterns. Because the ice is almost transparent, the background patch represents pure noise.

**Fig. 5.**
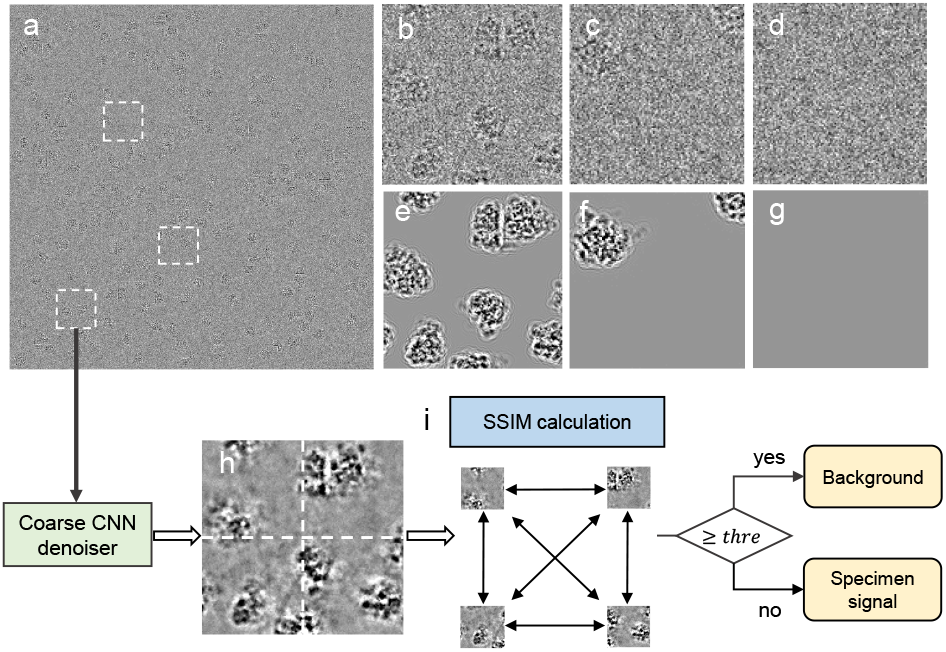
The illustration of the noise extraction algorithm based on patch similarity. (a) An illustration of the raw micrograph. (b-d) Three representative patches selected from the raw micrograph. (e-g) The clean corresponding specimen signals of the selected patches. (h) A patch enhanced by the coarse CNN denoiser trained with simulator; The patch is divided into *N* local patches with *N* = 4 in here. (i) The calculation of structural similarity (SSIM) and the determination of pure noise patches.

Here, we proposed a patch-similarity guided algorithm to extract the noise patches in a cryo-EM micrograph:

1. Given a micrograph *I*, denoise *I* with the CNN denoiser trained by the simulated data in subsection 3.2.1 to get an enhanced image *I*’;
2. Divide *I*’, into a set of overlapping patches Θ = {*P_i_*} (*d* × *d* pixel^2^ per patch) with a step size of *s*;
3. For each patch *P_i_* ∈ Θ, further divide *P_i_* into *N* local patches {*P_i,k_*} and calculate the structural similarity (SSIM) between each 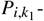 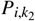 pairs (*k*_1_ ≠ *k*_2_);
4. For the patch *P_i_*, if ∀*k*_1_, *k*_2_(*k*_1_ ≠ *k*_2_), 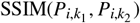 is large than a given threshold *thre*, determine *P_i_* as a background patch;
5. Repeat 1∼3 until all the background patches are identified, extract the exact patches in the original micrograph *I*.

It should be noted that the similarity determination is operated on micrograph *I*’, but the noise patch is extracted from the original micrograph *I*. The similarity measurement 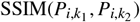 used in our algorithm is defined as

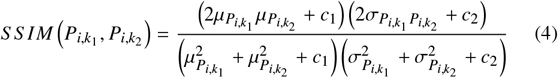

where 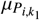 and 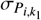 are the mean and standard deviation of 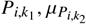 and 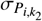 are the mean and standard deviation of 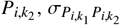 are the cross-covariance between patch 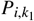 and 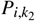, *c*_1_ and *c*_2_ are the constants that have very small values.

#### 3.2.3 Contrast-guided noise and signal re-weighting

Though we have designed an experimental parameters based simulation to produce simulated clean-noisy pairs (subsubsection 3.2.1), these simulated data is still not good enough to present the noise pattern in real cryo-EM images, which is more complex than the principles in the simulation. On the contrary, the GAN noise synthesizer (subsubsection 3.1.2) is able to produce noise patches with almost the same statistical properties as the real noise in the cryo-EM image. Here, we design a contrast-guided noise and signal re-weighted algorithm to transfer the noise pattern produced in the GAN noise synthesizer to the simulated clean data, to produce sophisticated clean-noisy pairs for fine CNN denoiser training.

Figure 6 shows the detailed process of the re-weighted algorithm. The algorithm accepts the pure noise patches {*V_i_*} generated from GAN and the clean signal patches {*S_i_*} generated from the simulation as input, utilizing the simulated patches {*X_i_*} corresponding to the exact signal patches from the simulation as a reference, and outputs a re-weighted dataset {*Y_i_*} with the GAN-synthesized noise transferred to the clean simulated signal. Here we use the contrast of *X_i_* as a baseline. Denote *Y_i_* = ℱ (*α* ∗ *S_i_* + *β* ∗ *V_i_* + *γ*) (ℱ(·) is a modulation function), the noise transfer should comply with the following objective function:

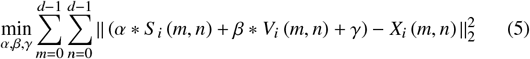

where *α*, *β* and *γ* are scalar coefficients and *d* is the patch size. Such a minimizing problem can be easily solved by the least-square method. Then, with the solved coefficients, the signal will be re-weighted and modulated, to produce clean-noisy pairs. The constructed clean-noisy pairs are fed into a fine CNN denoiser to train the model that captures the true noise statistics and restores specimen signal from noise.

**Fig. 6.**
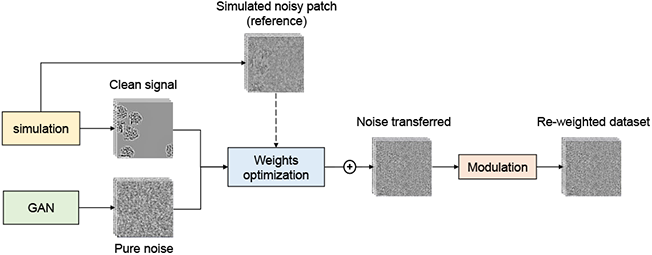
Contrast-guided noise and signal re-weighting. The noise pattern produced in the GAN noise synthesizer and the simulated clean signal are re-weighted with optimal coefficients and then modulated to produce clean-noisy pairs.

### 3.3 Implementation details

All the models are trained on two NVIDIA 2080Ti with 12GB VRAM.

The GAN noise synthesizer adopts a WGAN-GP network with the weights initialized from a standard Gaussian distribution with *σ* = 0.02. The slope of the leak of the LeakyReLU activation for both generator and discriminator is set to 0.2. Adam optimizer is used for model training with hyper-parameters set to 0.5, weight updating *β*_1_ = 0.999, and a learning rate of 0.0002.

The coarse CNN denoiser used for rough micrograph enhancement is trained with simulated datasets (subsubsection 3.2.1) with default initialization (Paszke *et al*., 2019). The fine CNN denoiser used for final denoising is trained with clean-noisy pairs generated by the GAN synthesizer and signal re-weight algorithm, initialized from the previous model weights. These two models are all trained using the Adagrad (Lydia *et al*., 2019) with a learning rate of 0.001. Given a trained denoiser, the denoising for a full-size micrograph is performed in overlapping patches. A padding approach is adopted to avoid the artifacts occurring at the patches’ edge.

## 4 EXPERIMENTS AND RESULTS

### 4.1 Datasets

Four simulated datasets and three real-world datasets are used to evaluate the performance of NT2C. The simulated datasets are generated with InsilicoTEM utilizing proteins downloaded from PDB and real-world datasets are collected from public repositories, including EMPIAR-10025 (abbr. EM25) (Campbell *et al*., 2015), EMPIAR-10028 (abbr. EM28) (Wong *et al*., 2014) and EMPIAR-10077 (abbr. EM77) (Fischer *et al*., 2016). The detailed information on these seven datasets are summarized in Table 1.

**Table 1.**
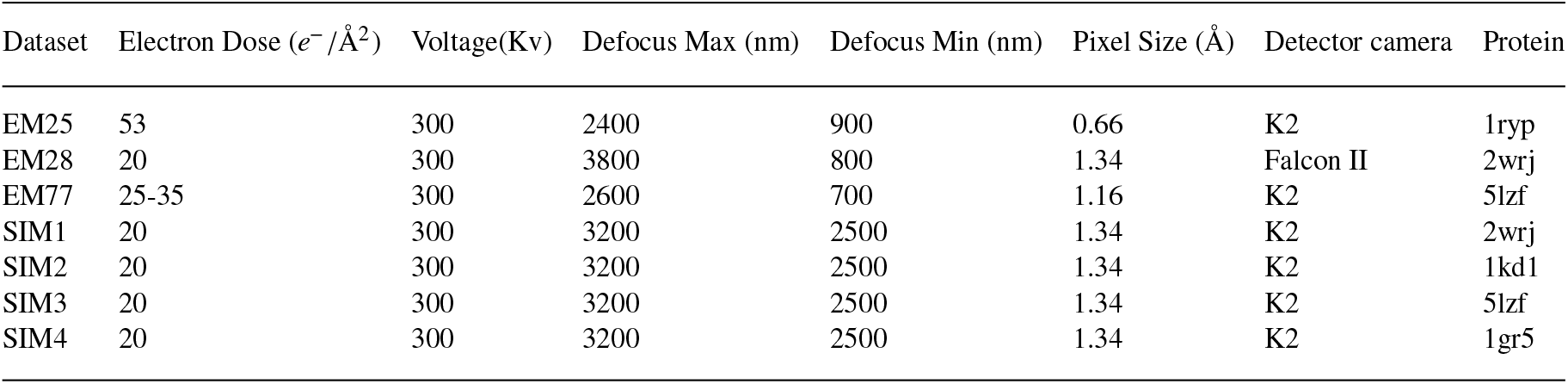
Three real-world datasets and four simulated datasets used for the denoisers training.

| Dataset | Electron Dose ($e^-/\text{\AA}^2$) | Voltage(Kv) | Defocus Max (nm) | Defocus Min (nm) | Pixel Size ($\text{\AA}$) | Detector camera | Protein |
| --- | --- | --- | --- | --- | --- | --- | --- |
| EM25 | 53 | 300 | 2400 | 900 | 0.66 | K2 | 1ryp |
| EM28 | 20 | 300 | 3800 | 800 | 1.34 | Falcon II | 2wrj |
| EM77 | 25-35 | 300 | 2600 | 700 | 1.16 | K2 | 5lzf |
| SIM1 | 20 | 300 | 3200 | 2500 | 1.34 | K2 | 2wrj |
| SIM2 | 20 | 300 | 3200 | 2500 | 1.34 | K2 | 1kd1 |
| SIM3 | 20 | 300 | 3200 | 2500 | 1.34 | K2 | 5lzf |
| SIM4 | 20 | 300 | 3200 | 2500 | 1.34 | K2 | 1gr5 |

### 4.2 Results

#### 4.2.1 Robustness of simulation-based noise pattern discovery

The simulation based on experimental parameters is the start point for coarse CNN denoiser training, which is critical to noise patch extraction and the consequent noise model discovery. However, the protein adopted in the simulation may be quite different from the unknown biological structures in real data.

To demonstrate the robustness of simulation-based noise pattern discovery, we generated four simulated datasets of protein 2wrj.pdb, 5lzf.pdb, 1kd1.pdb, 1gr5.pdb with the same simulation parameters and denoted them as SIM1, SIM2, SIM3, SIM4, respectively. Specifying SIM1 as the dataset to be denoised, we firstly trained three coarse CNN denoisers with SIM2-4 to roughly enhance the image contrast of SIM1. Then, we extracted the noise patches from SIM1 and learned the noise properties of these patches by a GAN noise synthesizer. The synthesized noise patches were then transferred to the clean signal patches of SIM2-4 for the training of fine CNN denoisers. The denoiser trained on clean-noisy pairs synthesized from xxxx.pdb is denoted as NT2C-xxxx. We tested these different denoisers on SIM1, whose results are shown in Figure 8. The NT2C-2wrj acts as a reference for comparison, which learns true noise patterns directly from clean-noisy pairs. It can be found that all these four denoisers have correctly resolved the specimen signal from the degenerated micrographs, resulting in a denoised output almost identical to each other.

**Fig. 7.**
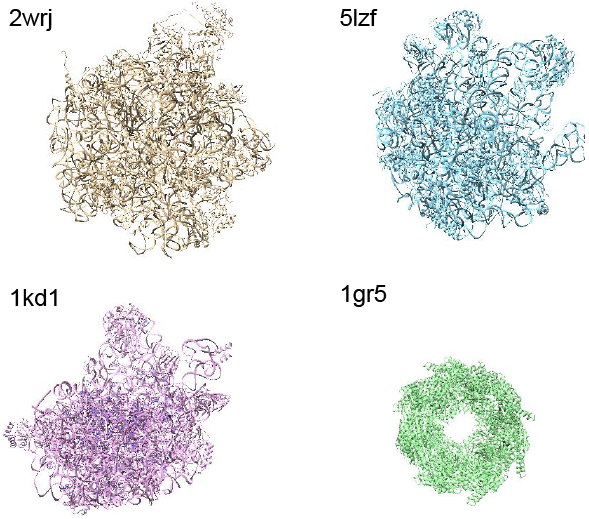
Illustration of the proteins used for simulated datasets generation and validation. PDB 2wrj is specified to generate the dataset to be denoised, while PDB 5lzf, 1kd1, and 1gr5 are the proteins used to generate peer simulation used in coarse CNN denoiser. Comparing with 2wrj, 5lzf has a similar structure and size, 1kd1 has a similar structure and smaller size, 1gr5 has a completely different structure and smaller size.

**Fig. 8.**
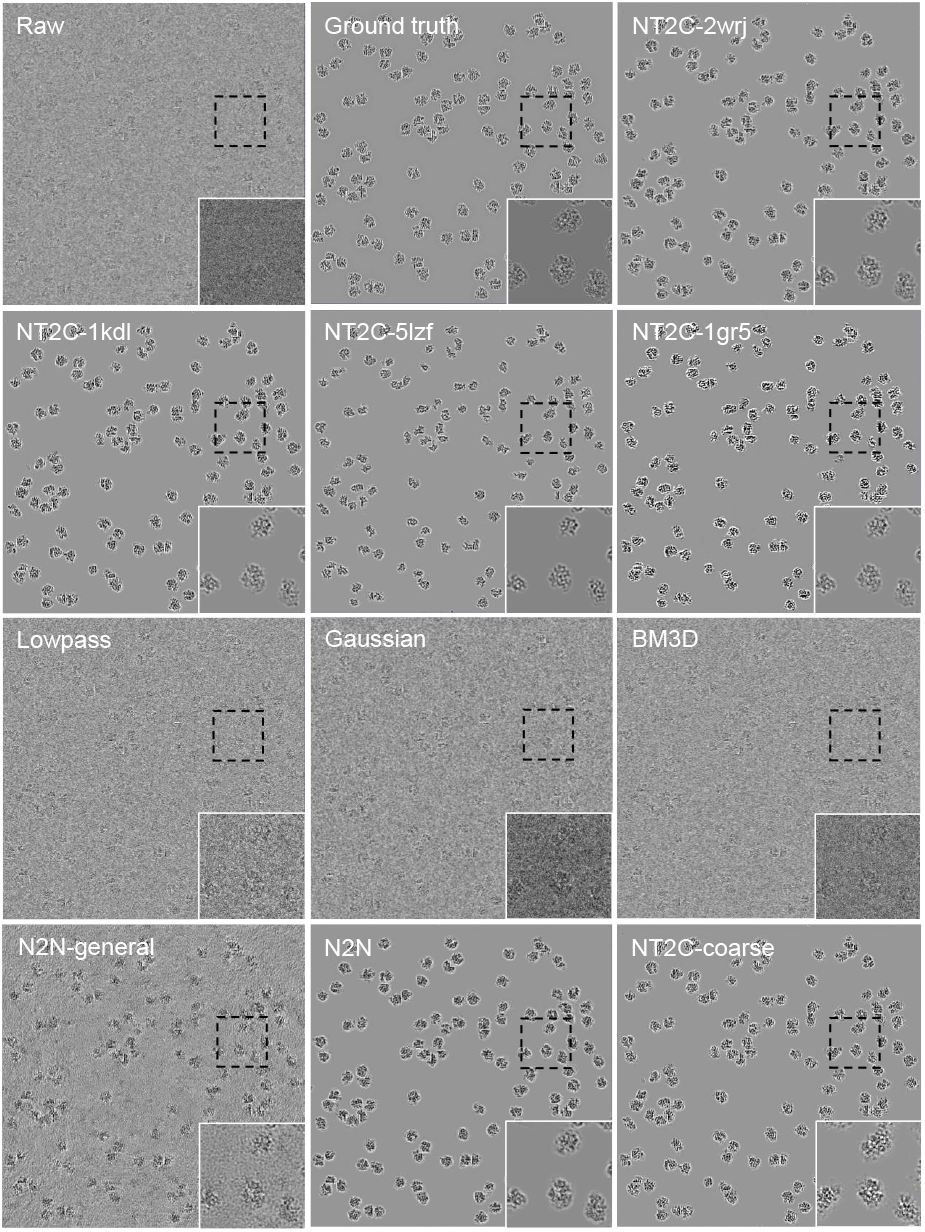
**1) Robustness of simulation-based noise pattern discovery**. The first two rows show raw micrograph, ground truth and denoised micrographs of denoisers trained with datasets simulated from 2wrj.pdb, 1kd1.pdb, 5lzf.pdb and 1gr5.pdb. **2) Denoising with NT2C improves micrograph interpretability and PSNR in simulated datasets.** The last two rows show comparison of different denoising methods on dataset generated with 2wrj.pdb, from left to right are Lowpass filter (2x binning), Gaussian filter, BM3D (third row), N2N-general, N2N and NT2C-coarse (last row). Detailed views of boxed region are shown at lower right corner of the micrograph.

Furthermore, with the availability of ground truth, we are able to quantitatively measure the denoising performance of different denoisers. The peak signal-to-noise ratio (PSNR) (Huynh-Thu *et al*., 2008), SSIM, and Pearson correlation coefficient (Pearson CC) (Benesty *et al*., 2009) between the denoised micrograph and the ground-truth were calculated and summarized in Table 2. It can be found that the size and structure of protein used in simulation hardly affected the noise pattern discovery and signal restoration of NT2C. Compared to NT2C-2wrj which reveals true noise distribution, the other three denoisers that trained with datasets simulated from different proteins achieved almost the same performance on all criteria.

**Table 2.** Comparison of four denoisers trained with different proteins based on SSIM, PSNR (in dB) and Pearson CC. (larger is better)

| Denoiser | SSIM | PSNR | Pearson CC |
| --- | --- | --- | --- |
| NT2C-2wrj | 0.846 | 29.78 | 0.823 |
| NT2C-5lzf | 0.842 | 29.78 | 0.823 |
| NT2C-1kd1 | 0.841 | 29.78 | 0.821 |
| NT2C-1gr5 | 0.843 | 29.77 | 0.822 |

##### Denoising with NT2C improves micrograph interpretability and PSNR in simulated datasets

We compared NT2C with four mainstream cryo-EM denoising methods, including three conventional methods, Low-pass filter, Gaussian filter (Haddad *et al*., 1991), BM3D, and a learning-based methods, N2N (Noise2Noise). Topaz-Denoise is employed for N2N where the general model provided by Topaz-Denoise is denoted as ‘N2Ngeneral’ and the Topaz-Denoise model retrained with a specific dataset is denoted as ‘N2N’. We also provide a coarse CNN denoiser that is used for roughly image contrast-enhancing in the noise extraction module of NT2C, denoted as NT2C-coarse. The comparison among different methods is carried on SIM1. Here, NT2C-1kd1 is adopted for comparison. The NT2C-coarse is trained with SIM2 that is generated with protein 1kd1.pdb.

The last two rows of Figure 8 present the denoised results of Low-pass filter (2x binning), Gaussian filter, BM3D, N2N-general, N2N, and coarse-NT2C. It can be found that both N2N and NT2C remove background noise while preserving the structural features significantly better than conventional methods. The very weak particle signals in the raw micrograph are clearly restored after denoised by N2N and NT2C. The image contrast is greatly enhanced by N2N-general. However, the noise is not completely removed. Especially, as shown in the small representative area, the image denoised by NT2C-coarse contains artifacts around the specimen signal, which do not appear in NT2C. NT2C can correctly capture the true noise distribution in SIM1 and remove it from the micrograph.

We further quantitatively assessed the performance of different methods based on the SSIM, Pearson CC and PSNR (in dB). As shown in Table 3, N2TC achieves the best performance on all three assessment criteria. Especially, our method improves SSIM by 0.05, Pearson CC by 0.06 and PSNR by *>*3 dB over N2TC-coarse which lacks true noise, and improves Pearson CC by 0.03 and PSNR by roughly 4 dB over N2N methods.

**Table 3.**
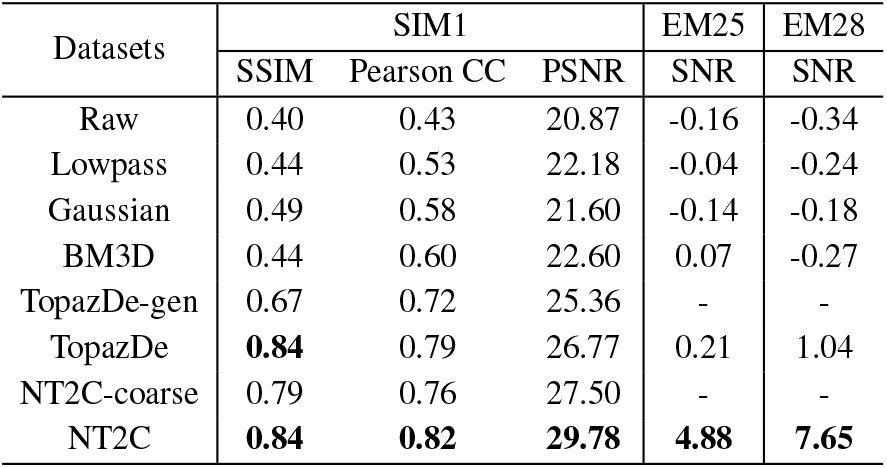
Comparison of denoising methods based on SSIM, Pearson CC, PSNR and estimated SNR (in dB, larger is better)

| Datasets | SIM1 |  |  | EM25 | EM28 |
| --- | --- | --- | --- | --- | --- |
|  | SSIM | Pearson CC | PSNR | SNR | SNR |
| Raw | 0.40 | 0.43 | 20.87 | -0.16 | -0.34 |
| Lowpass | 0.44 | 0.53 | 22.18 | -0.04 | -0.24 |
| Gaussian | 0.49 | 0.58 | 21.60 | -0.14 | -0.18 |
| BM3D | 0.44 | 0.60 | 22.60 | 0.07 | -0.27 |
| TopazDe-gen | 0.67 | 0.72 | 25.36 | - | - |
| TopazDe | **0.84** | 0.79 | 26.77 | 0.21 | 1.04 |
| NT2C-coarse | 0.79 | 0.76 | 27.50 | - | - |
| NT2C | **0.84** | **0.82** | **29.78** | **4.88** | **7.65** |

#### 4.2.2 Evaluation with Real Noise

To demonstrate NT2C’s ability to deal with complex latent noise in real cryo-EM images, we evaluated the performance of NT2C on three real-world datasets and compared it with Lowpass filter, Gaussian filter, BM3D and N2N.

##### Denoising with NT2C improves micrograph interpretability and SNR

Figure 9 shows a representative region selected from EM28 and denoised results of Low-pass filter (2x binning), Gaussian filter, BM3D, N2N, and NT2C. It can be seen that two learning-based methods, N2N and NT2C are better than conventional methods in noise smoothing and specimen signal enhancing. Moreover, NT2C achieves a stronger noise removal performance and reserved clearer specimen signal than N2N, therefore provides better interpretability.

**Fig. 9.**
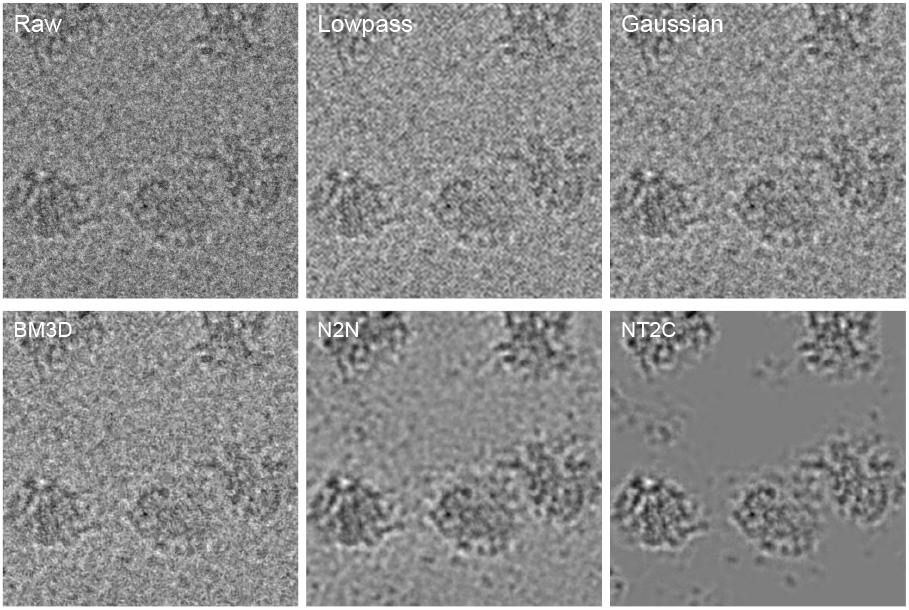
Denoising with NT2C improves micrograph interpretability in real-world datasets. Comparison among different denoising methods is carried out on real-world datasets EM28. A small region is selected to illustrates that NT2C performs better in noise smoothing and signal enhancing than both conventional (Lowpass filter, Gaussian filter, BM3D) and learning-based methods (N2N).

Furthermore, we quantitatively assessed denoising performance by measuring the SNR of raw micrographs and micrographs denoised with different methods. Due to the nonexistent ground truth, the SNR is estimated in a similar way to Bepler *et al*., 2020. Firstly, we select 10 paired signal and background regions across 10 micrographs where the background regions are as close as possible to the corresponding signal region. Given *N* signal and background pairs, 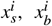 the mean and variance of each background region is marked as 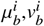 We define the signal for each region as 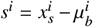 and calculate the mean and variance of signal region, 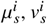. The average SNR in dB for the regions is defined as:

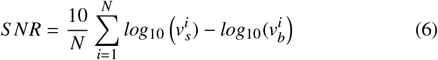

This SNR has no physical meaning, just criteria for comparison. As shown in Table 3, the conventional methods only improve roughly 0.1 dB over raw micrographs. NT2C method improves SNR by 8 dB over raw micrographs and roughly 6 dB over N2N methods.

##### NT2C accurately resolve complex noise model and restore clear specimen signal

To further study the NT2C’s performance on noise removal and specimen signal restoration, we selected two representative regions from dataset EM25, one containing specimen signals and the other containing pure noise. Figure 10 presents the raw micrograph and denoised results of Lowpass filter, Gaussian filter, BM3D, N2N, and NT2C. It can be found that the pure noise region denoised by NT2C is cleaner than all other methods, where N2N removes most of the noise and the conventional methods achieve the poorest performance on noise smoothing. Moreover, as shown in the region containing specimen signal, NT2C correctly decomplex structured features from complex noise. The signal restored by N2TC presents distinct structures and the particles with different projections are easily distinguished. Table 3 gives quantitative analysis on SNR, which further proves the notable performance achieved by NT2C. It improves SNR by 4.91 dB, 4.54 dB and >4.6 dB over raw micrograph, N2N and conventional methods.

**Fig. 10.**
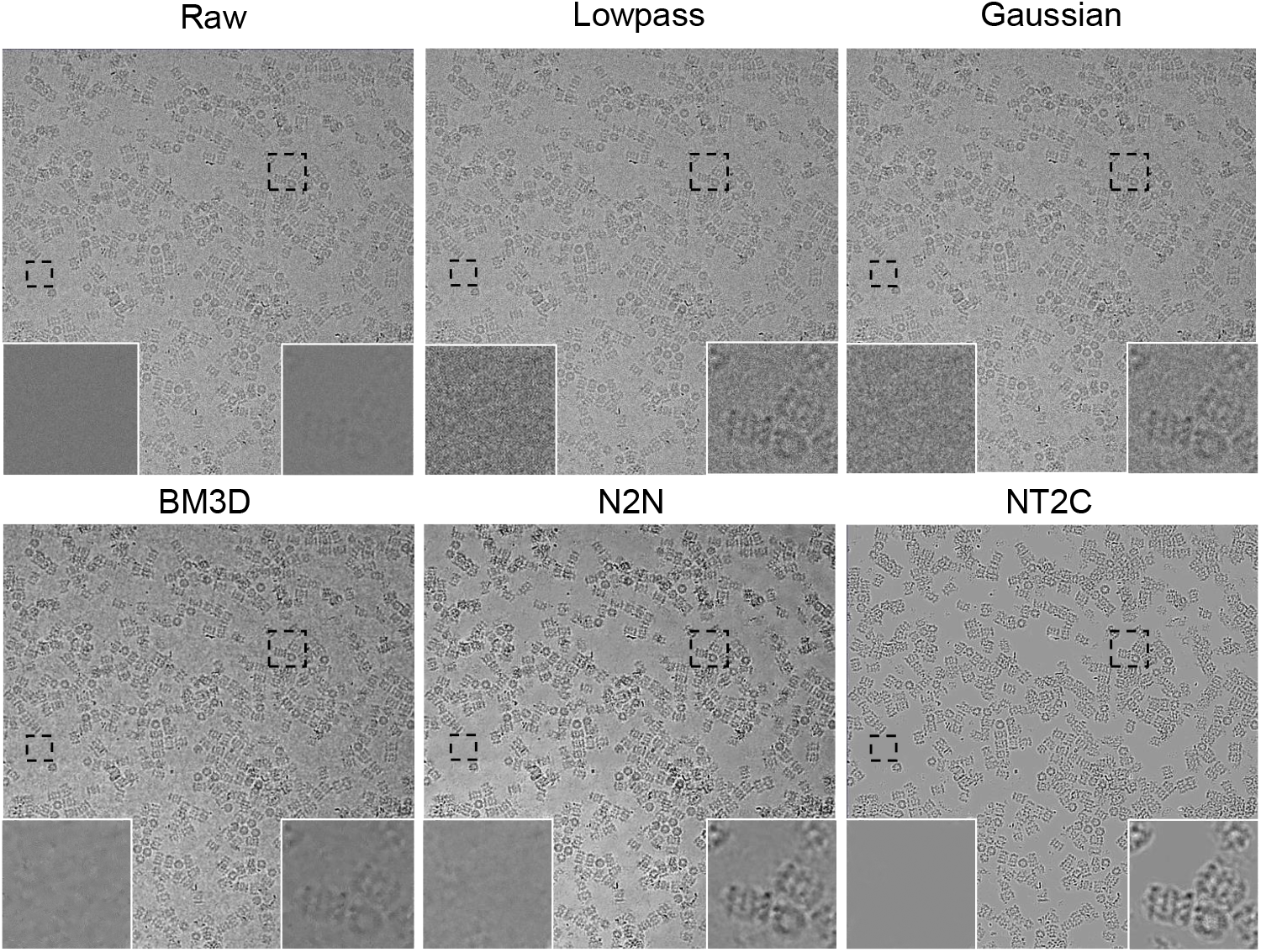
NT2C accurately resolve complex noise model and restore clear specimen signal. Two regions, one containing pure noise and the other containing specimen signal, are selected from one micrograph of EMP25 to illustrate the performance of NT2C. Comparing with Lowpass filter, Gaussian filter, BM3D, and N2N, NT2C can thoroughly remove noise and decomplex structured features from noise.

##### Denoising with NT2C enables more complete picking of hard-to-identify particles

We tested NT2C on micrographs with particularly difficult-to-identify particle projections, EM77, where the contrast of images is extremely low. Figure 11A shows representative micrographs before and after denoising. Before denoising, many particles were indistinguishable from noise by eye, such as the black-boxed regions a-c. After denoising, these particles in particular became readily identifiable.

**Fig. 11.**
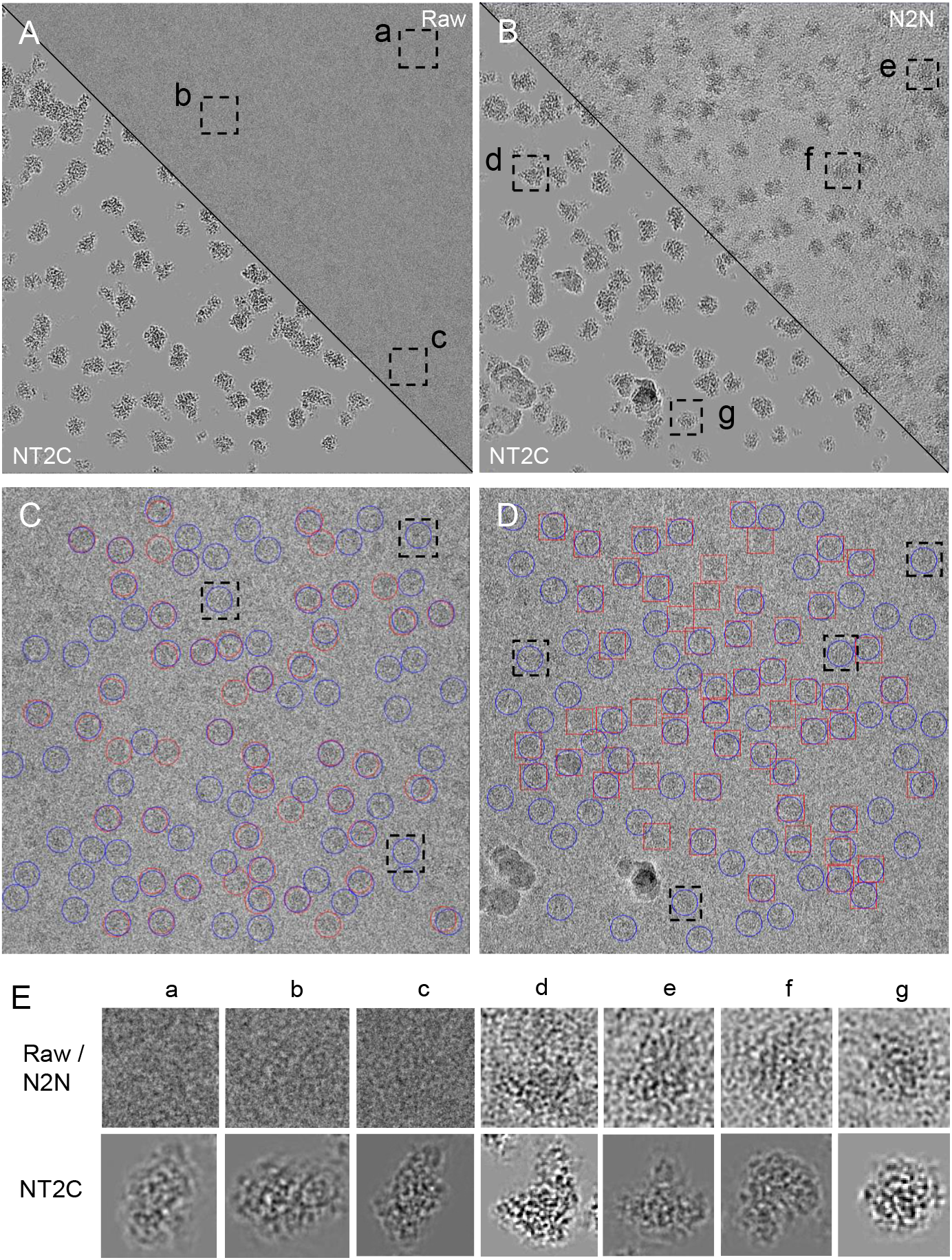
Denoising with NT2C enables more complete picking of difficult particle projections. (A) Micrograph from EM77 is split into the NT2C denoised and raw micrographs along the diagonal. (B) Micrograph from EM77 is split into the NT2C denoised and N2N denosied micrographs along the diagonal. (C) Particles picked by Topaz using raw micrograph (red circle) and micrograph denoised with NT2C (blue circle). (D) Particles picked by Topaz using micrograph denoised with N2N (red rectangle) and NT2C (blue circle). (E) The detailed views of particles boxed in black.

Since Topaz-Denoise is the most commonly-used denoising algorithm in cryoEM, we compared NT2C with Topaz-Denoise on improving particle picking. Figure 11B shows representative micrographs denoised by NT2C and Topaz-Denoise (denoted as N2N). Figure 11E d-g shows the particles that can be clearly recognized in micrograph denoised by NT2C. However, though the images contrast is enhanced by Topaz-Denoise, these particles are still difficult to indentify from background. NT2C greatly increases protein density confidence and allows researchers to identify low-density particle views from micrographs.

In order to quantify NT2C’s improvements on particle picking, we trained three Topaz (Bepler *et al*., 2019) particle denoising models with about 300 particles manually picked from raw micrographs and micrographs denoised by Topaz-Denoise and NT2C, marked as raw-picker, N2N-picker and NT2C-picker. The inference of NT2C-picker generates 45300 particles, N2N-picker generates 41998 particles, while raw-picker generates 38088 particles, where NT2C achieves the improvements on the recognition rate of hard-to-identify particles by roughly 8% over Topaz-Denoise and by 19% over raw micropgraphs. Figure 11C-D shows the particles picked from Figure 11A-B, where the particles from raw-picker, N2N-picker and NT2C-picker are denoted by red circles, red rectangles and blue circles. A lot of hard-to-identify particles, such as black-boxed regions, are recognized after the micrograph denoised by NT2C. The excellent denoising performance of NT2C will greatly relieve the difficulty of particle picking.

## 5 DISCUSSION AND CONCLUSION

In this article, we proposed a denoising framework for image contrast enhancement and specimen signal restoration in cryo-EM. The key idea of NT2C is discovering the noise model of cryo-EM images over pure noise patches and transferring the statistical nature of noise into the denoiser, making the denoising based on noise’s true properties. To cope with the complex noise model in cryo-EM, we further design a contrast-guided noise and signal re-weighted algorithm to achievie clean-noisy data synthesis and data augmentation for denoiser. To our knowledge, NT2C is the first denoising method that resolves the complex noise model in cryo-EM images. Comprehensive experiments on both simulated and real-world datasets demonstrate that NT2C is able to deal with high-level complex noise in cryo-EM images. A case study further demonstrates that NT2C can improve the recognition rate on hard-to-identify particles by 19% in the particle picking task.

Our follow work will further explore the complex relationship between image formation theory adopted in simulation and the real imaging process of cryo-EM, thus improving NT2C’s ability to restore information in real cryo-EM datasets.

## Notes

### Competing Interest Statement

The authors have declared no competing interest.

